# Parkinsonism disrupts neuronal modulation in the pre-supplementary motor area during movement preparation

**DOI:** 10.1101/2024.10.22.619634

**Authors:** Claudia M. Hendrix, Hannah E. Baker, Ying Yu, David D. Schneck, Jing Wang, Luke A. Johnson, Jerrold L. Vitek

**Affiliations:** Department of Neurology, University of Minnesota, Minneapolis, MN, 55455, USA; University of Minnesota Masonic Institute for the Developing Brain, Minneapolis, USA

## Abstract

Multiple studies suggest that Parkinson’s disease (PD) is associated with changes in neuronal activity throughout the basal ganglia-thalamocortical motor circuit. There are limited electrophysiological data, however, describing how parkinsonism impacts neuronal activity in the pre-supplementary motor area (pre-SMA), an area in medial frontal cortex involved in movement planning and motor control. In this study, single unit activity was recorded in the pre-SMA of two non-human primates during a visually cued reaching task in both the naive and parkinsonian state using the 1-methyl-4-phenyl-1,2,3,6-tetrahydropyridine (MPTP) model of parkinsonism. In the naive state neuronal discharge rates were dynamically modulated prior to the presentation of the instructional go-cue. In a subset of these modulated cells, the magnitude of modulation correlated linearly with reaction time (RT). In the parkinsonian state, however, modulation of discharge rates in the pre-SMA was disrupted and the predictive encoding of RT was significantly diminished. These findings add to our understanding of the role of pre-SMA in motor behavior and suggest that disrupted encoding in this cortical area contributes to the alteration of early preparatory and pre-movement processes that are present in Parkinson’s disease.

**SIGNIFICANCE STATEMENT:** Goal-directed movements, such as reaching for an object, necessitate temporal preparation and organization of information processing within the basal ganglia-thalamocortical motor network. Impaired movement in people with Parkinson’s disease is thought to be the result of pathophysiological activity disrupting information flow within this network. This work provides neurophysiological evidence linking altered motor preplanning processes encoded in pre-supplementary motor area (pre-SMA) neuronal firing to the pathogenesis of motor disturbances in parkinsonism.

## INTRODUCTION

Parkinson’s disease (PD) is a neurodegenerative disease associated with the death of dopaminergic (DA) neurons in the substantia nigra pars compacta (SNc) resulting in altered physiological activity throughout the basal ganglia-thalamocortical (BGTC) network (DeLong, 1990; Wang et al., 2017; Wichmann et al., 2017; Yu et al., 2021) and the development of parkinsonian motor signs (Galvan and Wichmann, 2008; Schnitzler and Gross, 2005; Vitek and Giroux, 2000). The supplementary motor complex (SMC), located in dorsomedial frontal cortex and consisting of the pre-supplementary motor area (pre-SMA) and SMA proper (SMAp), has strong connections to the BG (Watanabe et al., 2015) and is strongly influenced by the DA system (Albin et al., 1989; Alexander and Crutcher, 1990; Hikosaka and Isoda, 2010; Koirala et al., 2016; Koshimori et al., 2016; Watanabe et al., 2015). The SMC is involved in information processing and encoding of errors in behavior (Ballanger et al., 2007; Courtière et al., 2003; Pasquereau and Turner, 2013; Tomassini and Morrone, 2016), motor planning and temporal expectation (Cunnington et al., 2003; Hendrix et al., 2017; Leuthold and Jentzsch, 2001; Parker et al., 2020; Pasquereau and Turner, 2015; Tomassini and Morrone, 2016), and in organizing movements such as task switching, response selection and response inhibition (Muessgens et al., 2016). As a key nodal structure within the BGTC network strongly influenced by DA signaling, the pre-SMA has been implicated in the pathogenesis of motor disturbances found in parkinsonism (Nachev et al., 2008). This concept is further supported by multiple functional imaging studies have shown altered activity of the SMC in PD (Grafton, 2004; Playford et al., 1992). How parkinsonism impacts pre-SMA at the level of single neurons, however, remains unclear due to limited electrophysiological studies directly examining the changes in pre-SMA neuronal activity in the parkinsonian state during motor behavior.

We have previously shown that oscillatory neural activity derived from local field potentials (LFPs) within the SMC, especially within the pre-SMA, is disrupted in parkinsonism (Hendrix et al., 2017). The study by Hendrix et al. found that dynamic modulation of LFP activity correlated with subsequent reaction times in the healthy condition during a cued reaching task, but this predictive encoding of motor behavior was significantly diminished in the parkinsonian state. The goal of the current study was to expand upon these findings by examining the effect of parkinsonism on neuronal spiking activity within pre-SMA during pre-movement periods. We hypothesized that the neural mechanisms guiding the planning and preparation of movement within pre-SMA at the cellular level would be disrupted after the onset of parkinsonism. To test this hypothesis, we compared neuronal firing characteristics including patterns of modulation and neuronal encoding of reaction time in the pre-SMA of two nonhuman primates (NHPs) before and after induction of parkinsonism using the neurotoxin 1-methyl-4-phenyl-1,2,3,6-tetrahydropyridine (MPTP) while animals were engaged in a cued reaching task.

## MATERIALS AND METHODS

### Task and Surgical Procedures

All procedures were approved by the Institutional Animal Care and Use Committee of the University of Minnesota and complied with the United States Public Health Service policy on the humane care and use of laboratory animals. Two female NHPs (*Macaca mulatta;* T, 21 y.o and P, 18 y.o.) were trained to perform a cued reaching task (**Figure 1**) and implanted with a 32-channel electrode array targeting pre-SMA. The behavioral task and surgical approaches have been previously described (Connolly et al., 2015; Hendrix et al., 2013, 2017) and are briefly summarized below. During the reaching task, the animal placed its left hand on a start-pad to initiate trials and, after a variable delay, an instructional go-cue appeared on a touchscreen in 1-of-8 (NHP T) or 1-of-9 (NHP P) pseudo-randomized target positions. A successful reach to the target resulted in a juice reward and the animal initiated the next trial by voluntarily returning its hand to the start-pad. The time from trial initiation until the onset of the go-cue was defined as a preparatory preplanning time (PPT). Reaction time (RT) was defined as the time between the presentation of the ‘go-cue’ on the screen and reach initiation (i.e., hand leaving the start-pad). Reach duration was defined as the time between reach initiation and contact with the touchscreen monitor. The animal was required to maintain contact with the start-pad for the duration of the variable PPT (3-3.75 sec) and time limits were imposed on reach and reaction times (<0.8 sec). Failure to meet any of these criteria resulted in the immediate termination of the trial and exclusion from further analysis.

**Figure 1.**
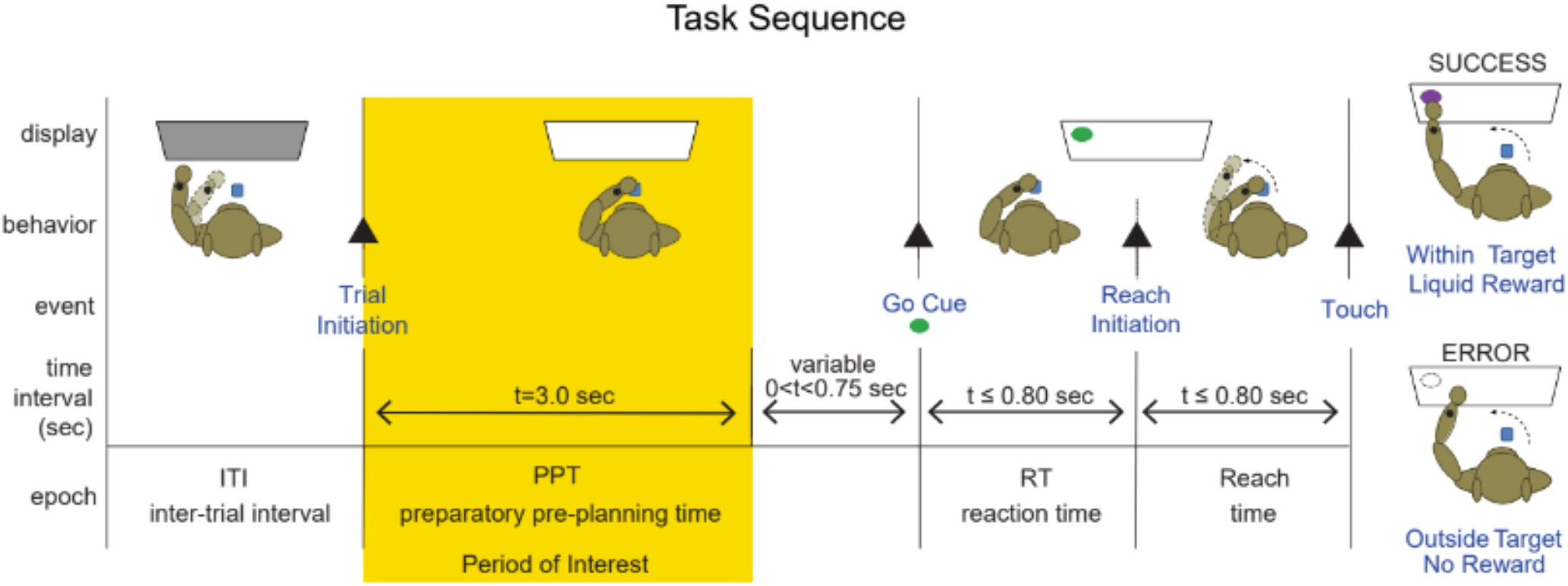
Task sequence and epochs of the visually cued center-out reaching task. The animal was seated in front of a touchscreen monitor (1st panel, display dark background) and start-pad (blue square), centered on the animal’s midline. Trials were self-initiated when the animal placed their hand on the start-pad and feedback of correct hand placement was confirmed by a change in touchscreen background color (2^nd^ panel, illustrated by display change from grey to white). The animal was required to maintain this initial holding position until the onset of a randomly presented go-cue (uniform random distribution between 3.0 and 3.75 s) or the trial was aborted. The go-cue appeared at one of eight (NHP T) or nine (NHP P) randomly presented target positions (3rd panel, 10 cm diameter, green circle). The animal was required to initiate and complete the reach within constrained time limits (RT < 0.8 s, reach < 0.8 s) or the trial was discontinued. Animals received immediate feedback indicating trial success (*top right*, touch within target, target color changes to magenta and juice reward) or error (*bottom right*, touch outside of target, target outlined, and no juice reward). The period of interest during which single unit activity was analyzed (yellow highlighted area) was defined as the first 3 s (t = 0–3 s) of the preparatory preplanning time (PPT).

Once stable task performance was achieved, the animal was implanted with a head-post and cephalic chamber positioned midline over the pre-SMA. After surgery, a microdrive (Gray Matter Research) with independently moveable tungsten microelectrodes (Alpha Omega, ∼0.3 MΩ, common reference grounded to the cranial chamber) was fixed to the cephalic chamber. Confirmation of the chamber placement and electrode positions relative to targeted structures were verified by co-registering preoperative MRI and postoperative CT scans using 3D slicer software package (Fedorov et al., 2012). Detailed methodology in cortical reconstruction and verification of electrode location has been previously published (Hendrix et al., 2017). Briefly, the CT imaging of the chamber wall and enclosed microdrive allowed for precise localization of each electrode position relative to cortical anatomical landmarks (**Figure 2A, B**). Cortical mappings of pre-SMA were reconstructed within each coronal slice using a standardized monkey atlas (Saleem and Logothetis, 2012) and scalable digital atlas (Bakker et al., 2015; Majka et al., 2012). The boundary between pre-SMA/SMAp was estimated based on published anatomical reconstructions (Akkal et al., 2007; Escola et al., 2003; Matsuzaka et al., 1992) using the coronal alignment of the 4 pre-SMA electrode locations with respect to the intersection of the upper/lower limbs and spur, or ‘genu’, of the arcuate sulcus.

**Figure 2.**
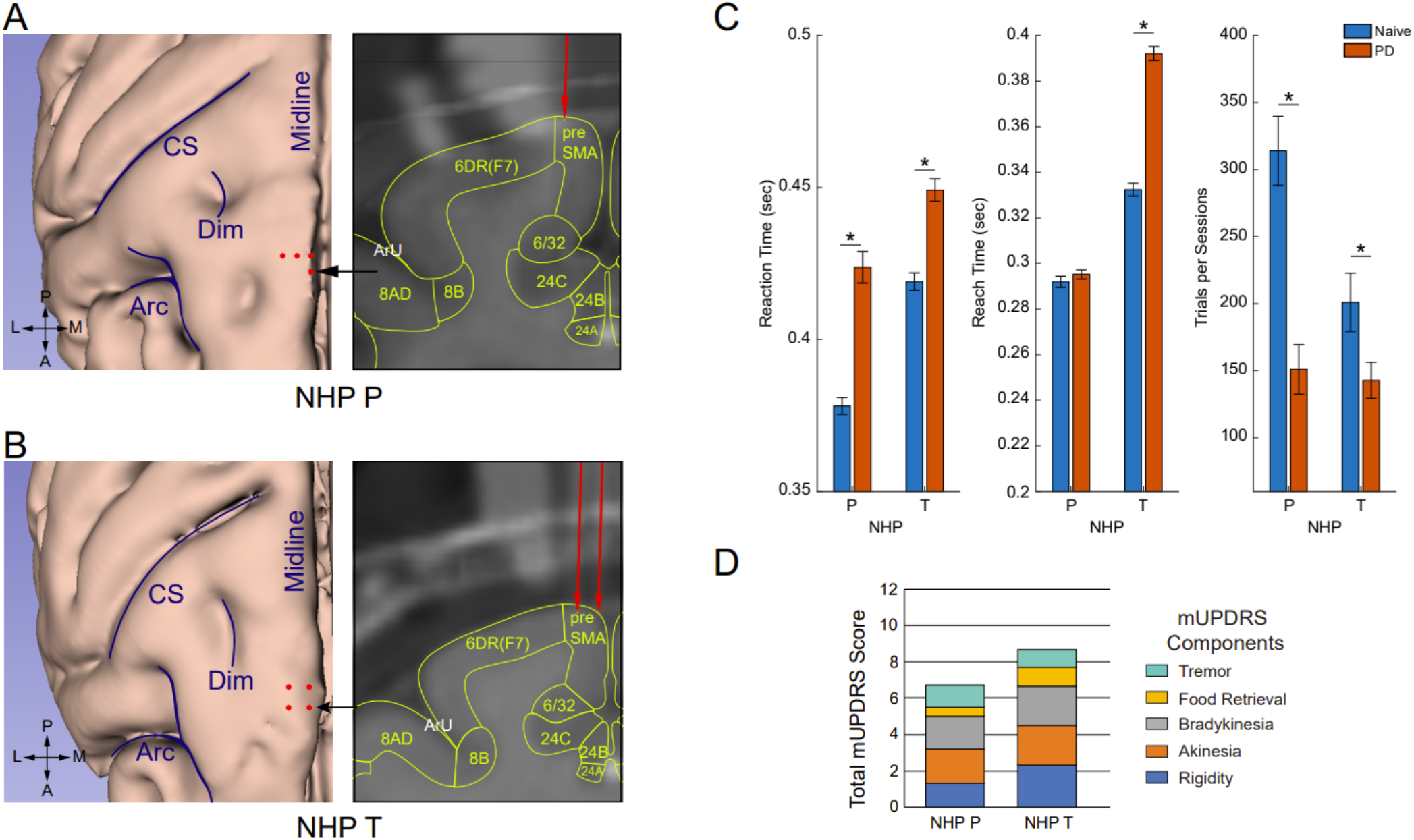
Reconstruction of cortical recording sites and behavioral changes after MPTP injections. **A-B**. Co-registration of preoperative MRI and postoperative CT were used to confirm electrode placements in animals P and T. *Left panels:* Custom 3D renderings of the animals’ cortex and relative placement of the pre-SMA electrodes with respect to cortical landmarks. *Right panels:* Coronal slice at the position of the most anterior electrode positions (black arrow) in the pre-SMA relative to adjacent cortical areas. CS - central sulcus, Dim - dimple, Arc - arcuate, ArU - uper limb of the arcuate. **C**. Behavioral performance in the naive (blue) and PD (red) state for each animal (mean and SEM). Reaction time (left), reach time (middle), and number of trials completed per session (right) are shown. *denotes significant (p<0.05) ANOVA F-value for RT / Reach Time and significant χ^2^ for Trails per Session. **D**. Clinical ratings (mUDPRS) of parkinsonian motor signs in each animal. Both animals were considered to be in a mild parkinsonian state (maximum mUPDRS total score possible = 27).

After data were collected in the naive state, the animals were rendered parkinsonian using the neurotoxin MPTP (Sigma Aldrich, 1 mg/ml solution). NHP T received a single intracarotid injection of MPTP (0.5 mg/kg). For animal P, systemic intramuscular injections were given weekly or biweekly to induce mild parkinsonism (cumulative total dose: 5.35 mg/kg). The severity of PD motor signs was assessed using a modified Unified Parkinson’s Disease Rating Scale (mUPDRS), which rated parkinsonian motor signs including akinesia, bradykinesia, rigidity, tremor for upper and lower limbs and food retrieval on the hemi-body contralateral to the site of neural recordings using a 0-3 scale (0=normal, 3=severe), with a maximum total score of 27 (Wang et al., 2022). Total scores were used to estimate overall motor sign severity (mild 1-9, moderate 10-17, and severe 18-27).

### SUA recording and post-processing

Neurophysiological data were recorded using a Tucker Davis Technologies (TDT) workstation operating at a sampling rate of 24.4 kHz. Raw data were bandpass filtered (300-5000 Hz) in Matlab (v2019b, Mathworks) and single units were isolated and sorted using principal component and template-based methods in Offline Sorter (Plexon). Further post-processing of the sorted units were performed using the Fieldtrip toolbox (Oostenveld et al., 2011) in Matlab.

Since we were interested in the neuronal mechanisms involved in early preparatory preplanning, the period of interest for SUA analysis was defined as the first 3.0 sec of the variable PPT (**Figure 1**, yellow highlighted area) during which the animal was in a holding-position. The firing rate modulation of each cell as well as correlations between changes in firing rates and behaviors across conditions were examined *on a trial-by-trial basis* to uncover patterns of neuronal activity that may otherwise be lost with conventional aggregate data analysis techniques. SUAs from each isolated unit were parsed into individual trials and time-locked to the onset of each PPT epoch (t = 0 sec). Trials in which movement related artifacts were found during the PPT were excluded from further analysis. Firing rates were estimated using a spike density function (Fieldtrip function: *ft_spikedensity*; inputs: time = 0.05 sec bins, frequency = 1 Hz bins).

### Firing rate modulation and correlation with reaction time

Two separate analyses were performed using cluster statistics: The first analysis determined if cell firing patterns were modulating during the PPT using a Monte Carlo method in which the significance probability (p-value estimate) is computed under a permutation distribution (n = 1,000). The Fieldtrip function *ft_timelockstatistics* was used (inputs: time = 0.05 sec bins, frequency = 1 Hz bins, method = ‘montecarlo’, α = 0.05, correctm = ‘cluster’, statistic=‘ft_statfun_actvsblT’) to generate an independent-samples t-statistic with a quantitative independent variable of mean baseline firing rate and a repeated measure variable of trial. Modulation was significant if the firing rate deviated from the mean for at least 150 continuous msec. Units without significant firing rate modulation were classified as tonic. Cluster-based nonparametric statistical inference testing in time series data has the advantage of controlling for i) the multiple comparison problem and ii) the sampling of time dependent observations, and validation of these methodologies have been extensively described (Maris and Oostenveld, 2007). Modulating cells were further described as having increasing, decreasing, or other firing patterns. Increasing and decreasing patterns were identified by taking the firing rate difference between the last and first second of the PPT using a paired two tailed t-test where significant p-values (p<0.05) indicated increasing or decreasing patterns of modulation and where non-significant p-values indicated other patterns.

The second analysis determined if on a trial-by-trial basis cell firing rates co-varied with subsequent RTs at any time during the PPT. We have previously detailed the use of this methodology when correlating LFP recordings and RT (Hendrix et al., 2017). Correlations between firing rates and RT used cluster-based nonparametric statistical inference testing. The Fieldtrip function *ft_timelockstatistics* (inputs: time = 0.05 sec bins, frequency = 1 Hz bins, method = ‘montecarlo’, α = 0.05, correctm=‘cluster’, statistic=‘ft_statfun_correlationT’) called for an independent-samples correlation rho-statistic with a quantitative independent variable of RT and a repeated measure of trial. A resulting temporal mapping of correlation values (rho) between RTs and firing rate bins (time-rate) was obtained.

### Population statistics

Population statistics were used to assess the proportion of cells with significant firing rate modulation or correlation with RT at any given time point in the PPT analysis window. The population statistic used frequency counts, the number of significant observations relative to the total, to determine the percentage of total neurons. An array of binary values (0 or 1) was assigned to each time bin indicating the absence or presence of significant t-values and correlation values, respectively, as determined by each cluster analysis (see above). The summation of binary arrays provided overall counts of SUA recordings with significant values at each time bin. Contingency analysis of significant counts against total numbers was used to compute the Likelihood-Ratio Chi-square (χ^2^) for each condition and related probabilities. Unless otherwise indicated a threshold of significance for all inference testing was p<0.05.

### Bootstrapping statistics

To test for temporal differences in the percentage of cells with significant firing rate modulation across naive and PD conditions we used the bootstrap technique, which is a robust nonparametric procedure that uses resamples of the data (Efron and Tibshirani, 1994). To ensure numerical precision 10,000 bootstrapped samples were used to create 95% confidence bands (Hogan and Laird, 1996). A 95% confidence band was constructed for the difference between the percentage curves from the naïve and PD conditions. The time points for which the confidence band for the difference did not contain zero meant that the difference between the modulation and encoding curves was statistically significant at those time points. We also were interested in whether there was a difference between naïve and PD states in encoding over the time course of the PPT. To address this question, we examined the first and last 1-sec segments of the PPT by constructing 95% bootstrap confidence intervals (CI) for the difference in encoding, where the lower and upper bounds of the CI were calculated from the standard deviation of the bootstrapped mean differences. CIs that did not contain zero indicate significant difference in encoding.

### Behavior Data Analysis

Objective measures of task performance including the RT, reach time, and trials completed were calculated. Differences in RT and reach were tested using a repeated-measures mixed-model nested factorial design model with a fixed effect (condition, naïve-PD) and random effect (recording session) nested within the condition. Differences in the number of trials completed per session were tested across conditions using a nonparametric Wilcoxon rank order test.

## RESULTS

### Summary of behavioral differences and severity of parkinsonian signs

Behavioral performance in the touchscreen task was impaired in the parkinsonian state in both animals (**Figure 2C**). Specifically, reaction times were significantly shorter in both animals (animal P, F(1,288)=67, p<0001; animal T, F(1,359)=41, p<0.0001), reach time durations significantly increased for animal T (animal P, F(1,288)=1, p=0.32; animal T, F(1,359)=42, p<0.0001), and fewer trials were completed per recording session in both animals (animal P, F(1,68)=27, p=<0.0001; animal T, F(1,49)=6, p=0.02). Post-MPTP clinical ratings averaged across sessions indicated that the animals were mildly parkinsonian with mUPDRS total scores (± standard deviation) of 6.6 ± 0.2 and 8.7 ± 0.7 for animals P and T, respectively (**Figure 2D**).

### Pre-SMA firing rate modulation is reduced in parkinsonian state

A total of 591 cells collected from 2 animals were included in the study (see **Table 1**). In the naïve state, in both animals the majority of pre-SMA cells modulated their firing rates during the PPT period of the task. An example of a cell classified as modulating is shown in **Figure 3A**. Increasing firing rates over time can be observed in both the aggregate average over all trials and in the spike activity across individual trials. In this example, the result of the trial-based cluster analysis indicated that the firing rate during the first ∼1.4 sec of the PPT was significantly lower than the average firing rate followed by a period starting at ∼1.8 sec in which the rate was higher than the average firing rate. An illustrative example of a cell classified as tonic with its firing rate remained relatively constant over the duration of the PPT is shown in **Figure 3B**. In this case the result of the trial-based cluster analysis did not detect significant deviations in firing rates from the mean. The percentage of cells classified as modulating decreased significantly after MPTP administration resulting in a population of predominately tonically firing neurons in the pre-SMA in the parkinsonian state (**Figure 3D;** animal P, χ^2^(1,230)=78, p<0.0001; animal T χ^2^(1,361)=96, p<0.0001)

**Table 1.** Summary of recording sessions and cell counts.

| NHP ID | Sessions |  | Total Cell Counts |  | Tonic Cells |  | Modulating Cells |  | RT Encoded Cells |  |
| --- | --- | --- | --- | --- | --- | --- | --- | --- | --- | --- |
|  | Naive | PD | Naive | PD | Naive | PD | Naive | PD | Naive | PD |
| P | 25 | 45 | 127 | 103 | 23 | 77 | 104 | 26 | 14 | 0 |
| T | 23 | 28 | 188 | 173 | 62 | 143 | 126 | 30 | 35 | 9 |

**Figure 3.**
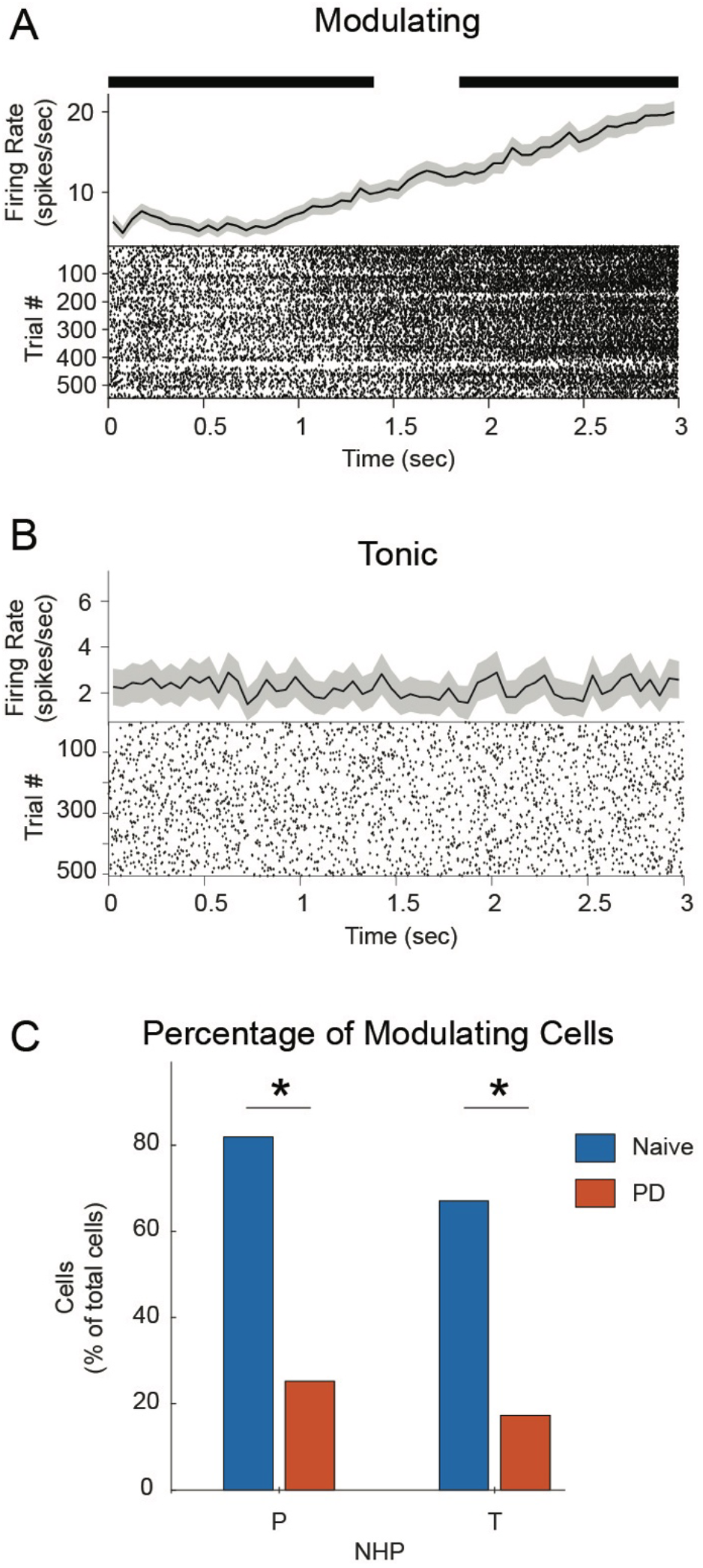
Loss of modulation in pre-SMA in parkinsonism **A**. Illustrative example of a modulating cell with a firing rate that deviated relative to the mean firing rate during the PPT. Raster plot of spike times across trials is shown, along with average firing rate and 95% CI across trials. Black bars indicate significant deviation of the firing rate relative to the mean. **B**. Illustrative example of a cell with a tonic firing rate (same nomenclature as above) where there was no significant period(s) of deviation from the mean. **C**. Bar plots show that in both animals the percentage of the total cells classified as modulating significantly decreased from naïve (blue) to PD (red) state (*p<0.05, χ2).

Modulating cells were further categorized according to their firing rate changes during the PPT (**Figure 4A**). Representative examples are shown for cells with increased, decreased, and other patterns in firing rates. A breakdown of the percentage of all modulating cells by pattern is summarized in pie charts (**Figure 4B**). In the naïve state, most modulating cells were either increasing or decreasing in firing rate during the PPT. In the parkinsonian state the majority of cells fell into the “other” category, though this change in proportion only reached significance in animal T.

**Figure 4.**
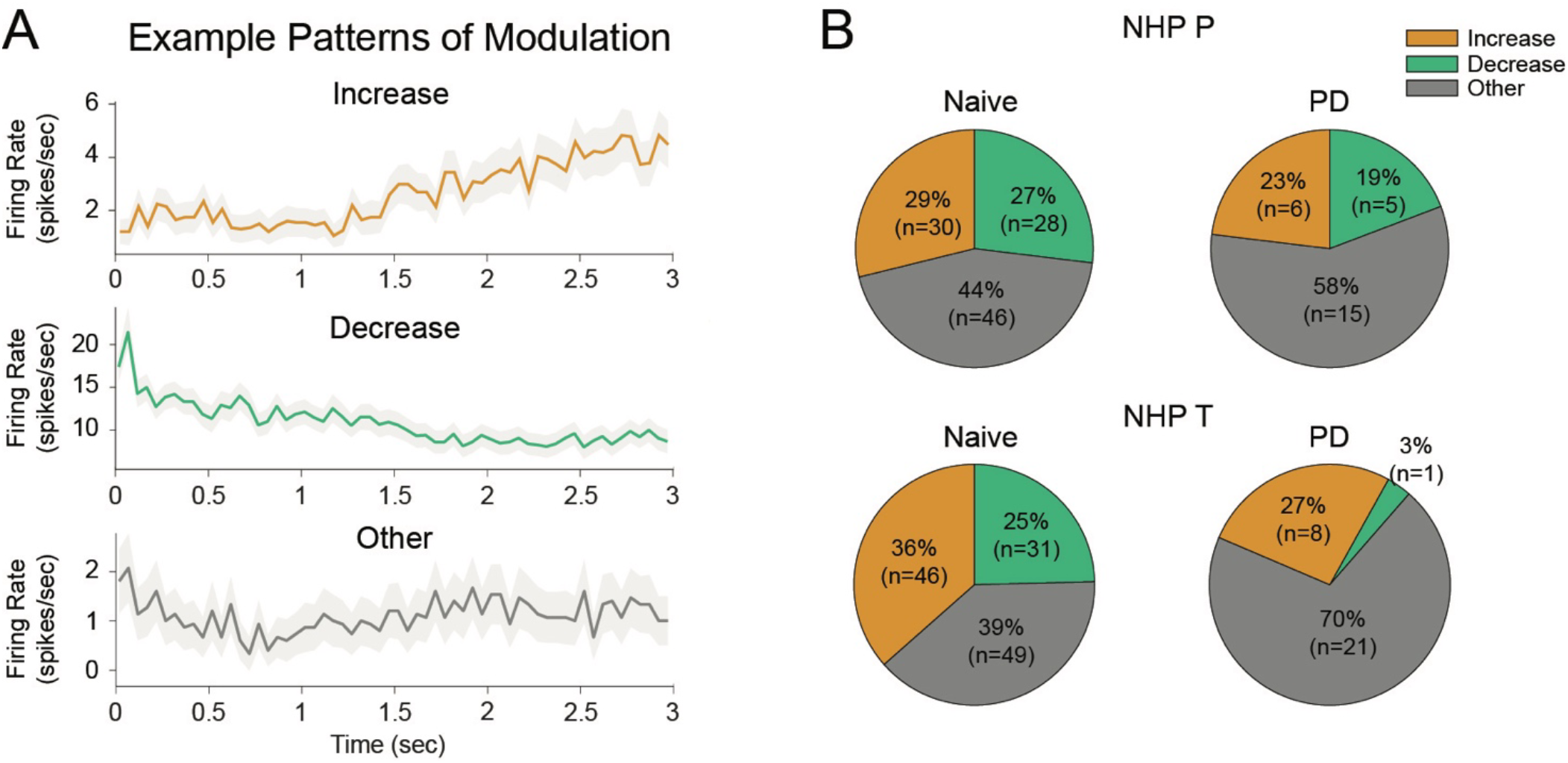
**A**. Modulating cells were classified as having increasing (top), decreasing (middle), or other (bottom) firing rates during the PPT. Each line plot illustrates the mean (solid) and CI of the mean (shaded) firing rate over the duration of the PPT. **B**. Breakdown of the percentage of total modulating cells by pattern in the Naive and PD states (left and right columns, respectively) in animal P and T (top and bottom row, respectively).

Having determined that fewer cells in the parkinsonian state had significant firing rate modulation during the PPT we then examined the temporal characteristics of firing rate modulation across the population of recorded cells in each state, as illustrated in **Figure 5A** (NHP P, all modulating cells in the naive state). In animal P for example, cells with significant firing rate modulation in the naive state tended to cluster within the early and later periods of the PPT. These observed temporal characteristics are summarized as a percentage of total cells with significant modulation during the PPT (**Figure 5B**). Consistent with the population statistics showing that the percentage of modulating cells decreased in parkinsonism, for both animals the curves of the percentage of modulating cells over time shows an overall decrease in the PD state. Specifically for animal P, a significantly larger percentage of the total cells were modulated during the early (∼0.1-0.6 sec) and later (∼2.3-3 sec) period of the PPT in the naïve state compared to the parkinsonian state (**Figure 5B**, left panel, gray bars). Similar findings were noted in animal T (**Figure 5B**, right panel, gray bars) where modulating cells in the naive condition preferentially modulated during the early PPT (∼0.1-0.6 sec) and this preferential period of modulation was lost in PD.

**Figure 5.**
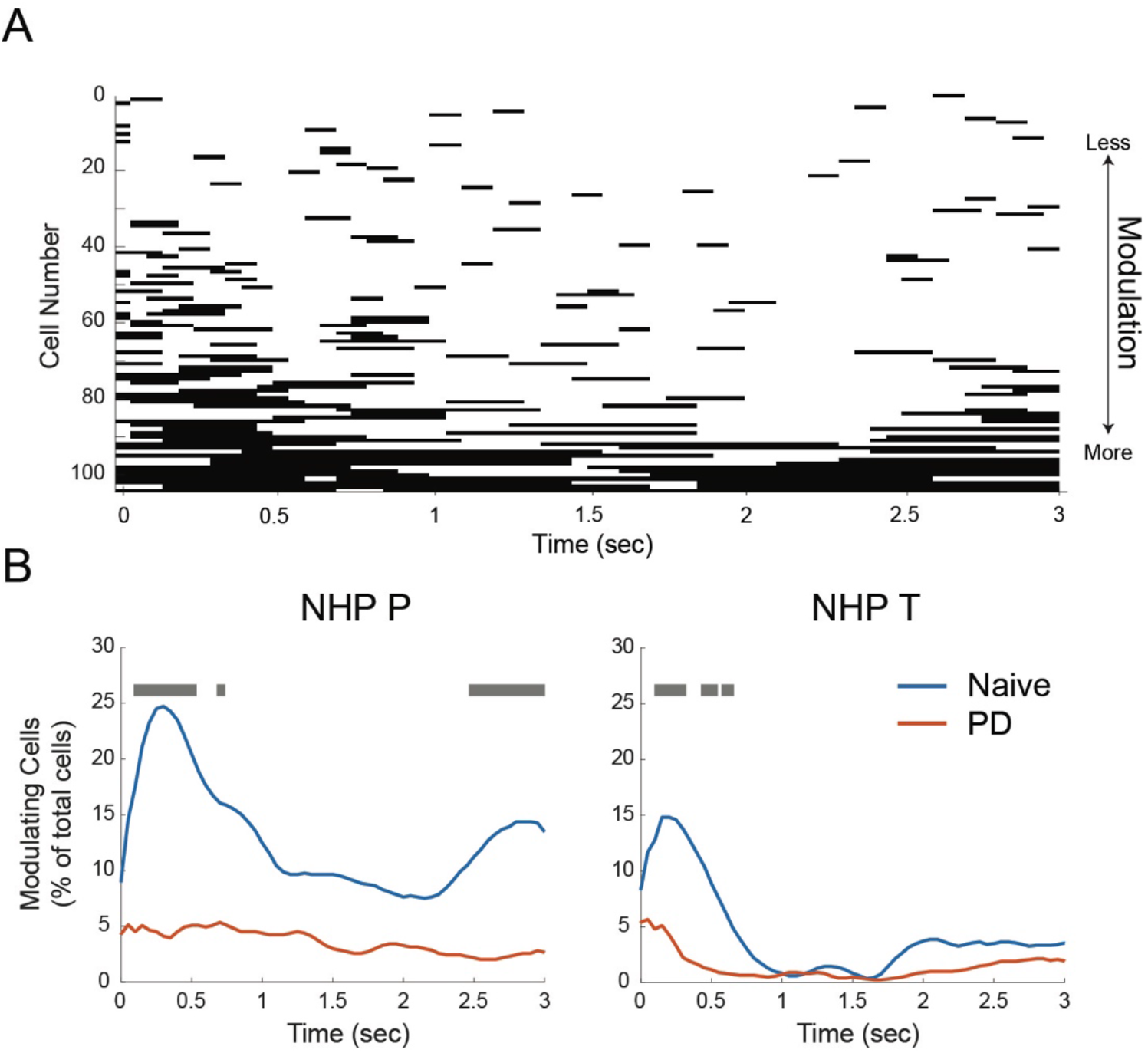
Temporal changes in modulating firing rates were lost in PD. **A**. Periods of significant change in firing rates relative to the mean firing rate by cell (cell number) during the PPT (t=0-3 sec) for all cells recorded in animal P in the naive state. Each cell (row) had periods of significant firing rates that deviated from the mean firing during the 3 sec PPT. Cells are sorted by degree of modulation (less to more in ascending order from top to bottom). in the naive (blue bars) and PD (red bars) states **B**. The percentage of total cells with significant modulation during each time bin (50 msec bins) in the naive (blue line) and PD (red line) for animal P (left panel) and T (right panel). Gray bars indicate time bins where significant differences were found between the Naïve (blue) and PD (red) traces. **C**. Breakdown of the percentage of total modulating cells by pattern in the Naive and PD states (left and right columns, respectively) in animal P and T (top and bottom row, respectively).

### Correlations of pre-SMA neuronal firing with reaction time

A subset of pre-SMA cells had firing rates during the PPT that predictively encoded reaction times. A representative example illustrates, for a single cell recording in animal P in the naive state, that the magnitude of the cell firing rate for each trial correlated with subsequent reaction times (**Figure 6A**). In this example, significant negative correlations between the firing rate and the RT were found towards the later part of the PPT (2.2-3 sec). This correlation analysis was carried out for all recorded cells, however only cells that had been classified as modulating were found to encode RT. In both animals, the percentage of cells with predictive encoding significantly decreased in the parkinsonian state (animal P, χ^2^(1,230)= 17, p<0.0001; animal T χ^2^(1,361)=16, p<0.0001) (**Figure. 6B**). The temporal characteristics of the population cells with predictive encoding during the PPT as a percentage of total cells over time is shown in **Figure 6C**. In the naïve state in both animals there was a trend towards increased percentage of cells encoding of reaction time towards the end of the PPT, reaching significance in animal T (mean diff=0.018 and 95% bootstrap CI = 0.002, 0.032). This trend was not observed in the parkinsonian state in animal T (mean diff =0.0005 and CI = -0.004, 0.0195), and no cells in the parkinsonian state in animal P encoded reaction time.

**Figure 6.**
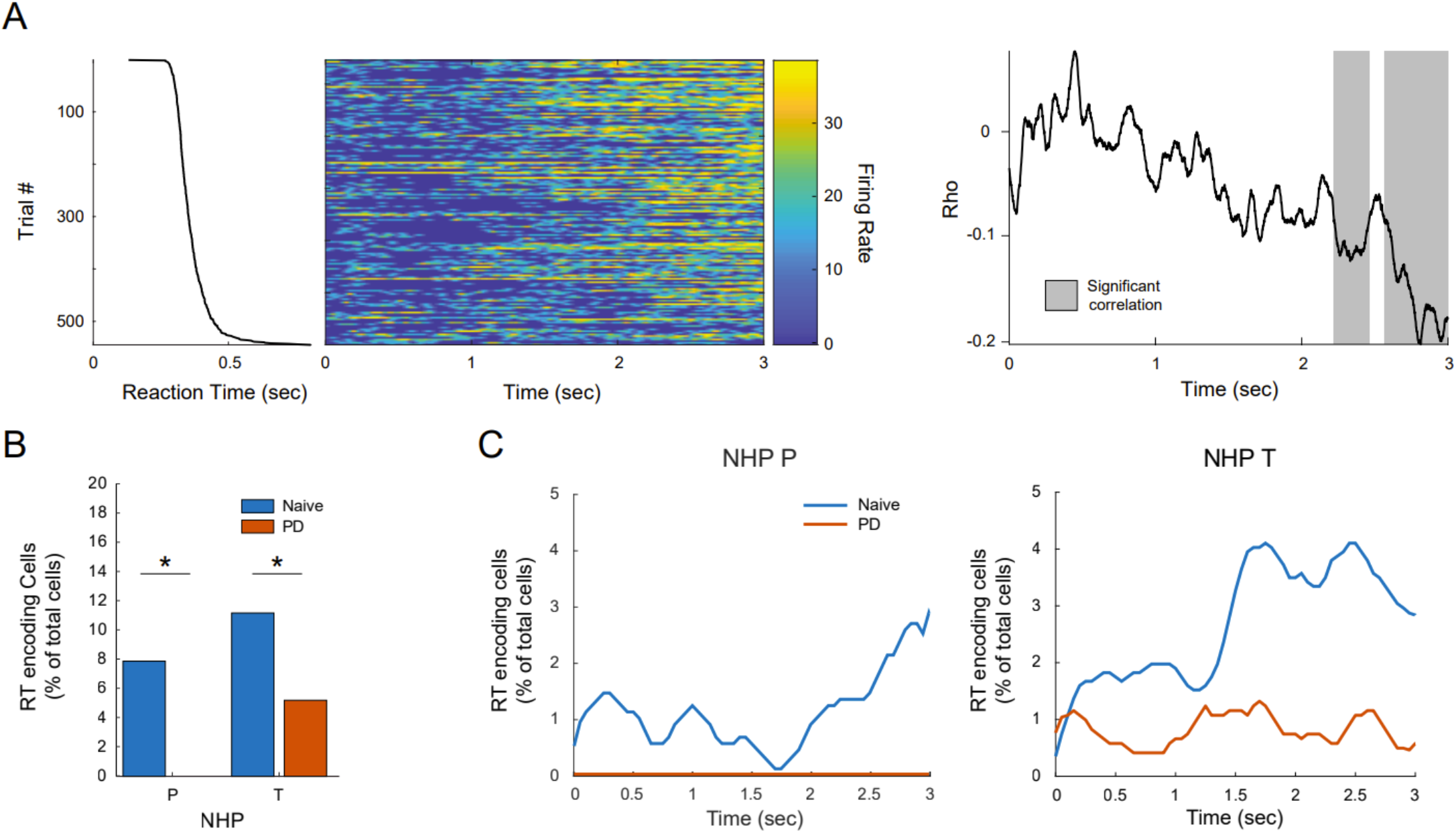
Pre-SMA cell encoding of reaction time is diminished in the PD state. **A**. Illustrative example of a cell with a firing rate during the PPT that correlated with subsequent reaction times. Trial reaction times sorted by duration (ascending, top-to-bottom) during a recording session in the naive state. Color maps of pre-SMA SUA firing rates by trial (each row) time-locked to onset of the PPT (t=0) and sorted by reaction time. Statistical mapping of correlation values (rho) between the reaction times and binned firing rate (50 msec) where clusters (grouped-bins) of significant correlation values are highlighted (gray shaded areas denote significant time bins). **B**. Barplots show the percentage of modulating cells with firing rates that correlated to reaction times in the naive (blue bars) and PD (red bars) states significantly decreased ‘* p<0.05 χ^2^ test in both animals with onset of parkinsonism. **C**. The percentage of all cells with significant correlations between the cell firing rates and reaction time during each time bin (50 msec bins) in the naive (blue) and PD (red) for animal P (left panel) and T (right panel). In the PD state, no cells encoded reaction times in animal P (flat red line at 0%).

## DISCUSSION

In this study we investigated the effects of MPTP-induced parkinsonism on neuronal activity in the pre-SMA during a “holding state” preceding cued movement in two NHPs. A distinct advantage of this preparation not feasible in human studies is that it allowed for a within-subject comparison of changes in activity of neuronal cell populations between healthy and parkinsonian conditions. A strength of our trial-by-trial analysis approach was that we were able to examine the relationship between pre-SMA neuronal firing and motor behavior (i.e., reaction time). Induction of the parkinsonian state was associated with 1) a dramatic reduction in the percentage of pre-SMA neurons with firing rate modulation during the holding period and 2) diminished pre-SMA neuronal encoding of reaction time. These findings add to our understanding of the role of pre-SMA in motor behavior and suggest a fundamental role of this cortical area in early preparatory and pre-movement processes that are altered in parkinsonism.

### Pre-SMA firing rate modulation in the naïve state

The present study focused on a period of the task often referred to as a baseline period (i.e., a waiting/holding period of the task before cued instruction is provided) during which neural activity is often assumed stationary. We found, however, that a large proportion of cells in the pre-SMA in healthy animals had significant firing rate modulation during this period. Most modulating cells had a gradual increase or decrease in firing rate during the PPT. Similar pre-movement pre-SMA activation patterns were found in a previous study in naïve NHPs and referred to as “buildup” activity (Matsuzaka et al., 1992). In their study, animals were required to press one of two targets according to a visual cue after a variable delay period. The authors reported that the majority of pre-SMA cells had firing rate modulation during this delay period, suggestive of involvement in movement preparation, but they did not provide further explanation of the functional significance of these “buildup” cell types. One might speculate that the buildup activity noted in their study is related to the processing of instructional cues that were provided before the delay period during the task. Task parameters in the current study differed, however, in that the animals were not provided with target-specific information before or during the pre-movement waiting period. Yet we found similar buildup activity in the majority of modulating cells in the naïve state. Thus, the pre-movement buildup patterns in the pre-SMA observed in both studies may reflect, in part, anticipation of and preparation for cued movement onset more generally rather than target-specific movement planning.

### Reduced Pre-SMA modulation in the PD state

In the present study, after the induction of the PD state the percentage of cells classified as modulating decreased significantly resulting in predominately tonic firing rates across the cell population during the PPT. Moreover, the preferred timing of significant modulation during the early and later PPT is replaced with non-specific other patterns of modulation that are lacking the temporal coordination present in the naïve animals (**Fig. 5B**). It has been suggested the DA loss in parkinsonism may induce hypoactivation in pre-SMA activity via reduced mesocortical dopamine signaling to medial frontal cortex (Escola et al., 2003). Our finding of a significant decrease in the percentage of the cell population identified as modulating would be consistent with this concept of pre-SMA hypoactivation. Multiple functional imaging studies have also reported hypoactivation of the pre-SMA in patients with PD compared to healthy controls (Mallol et al., 2007; Sabatini et al., 2000). There are several additional mechanisms that could potentially underly changes in pre-SMA activity that are observed in PD. It has been suggested that cerebellar inputs via the thalamus provide anticipatory activation of the supplementary motor complex (Rahimpour et al., 2021), and there is increasing evidence that cerebellar processing is disrupted in PD (for review see (Wu and Hallett, 2013)). It may also be the case that structural changes within the pre-SMA contribute to the observed changes in neuronal activity. A histological study comparing PD patients to healthy controls found a selective loss of cortico-cortical projecting pyramidal neurons in the pre-SMA in PD (MacDonald and Halliday, 2002). The authors suggest that the pathogenesis of PD includes a degeneration of premotor projections from the pre-SMA in addition to basal ganglia dysfunction. Together these studies combined with our findings support the idea that disruption in pre-SMA structure and function contributes to altered movement planning processes that are prominent in PD (Fasano et al., 2022; Gentilucci and Negrotti, 1999).

### Predictive encoding of reaction time is disrupted in PD

The neurophysiological finding of fewer pre-SMA cells with significant modulation combined with the behavioral finding of slowed average reaction times in the PD state motivated us to further investigate whether a direct relationship between neuronal firing and task behavior could be identified. Addressing this question necessitated a trial-based analysis approach to examine the relationship between each pre-SMA neuron firing responses during the PPT and subsequent reaction times. Indeed, we found that in the naïve state a subset of neurons had firing rates that significantly correlated with RT. This predictive encoding of behavior was significantly diminished in the PD state. Similarly, we have previously shown that early preparatory and pre-movement processes in pre-SMA LFPs are altered in parkinsonism (Hendrix et al., 2017). Specifically, beta band modulation normally correlated with subsequent RTs is lost in PD. This study provides the first trial-by-trial neurophysiological evidence linking altered motor pre-planning processes encoded in pre-SMA single unit activity to impaired motor behavior in PD. Together these two studies point to a disruption of pre-SMA predictive encoding as a potential cortical mechanism underlying prolonged RT and errors in executive function that are present in PD. Future studies specifically examining concurrent spike-LFP modulation in relation to preparatory behaviors may further elucidate the functional significance of LFP and neuronal population changes in the pre-SMA to the manifestation of PD motor signs.

### Limitations and Future Directions

The present study focused on a single cortical area. Future studies are needed to determine whether the loss in predictive encoding we report originates within the pre-SMA or within brain regions projecting to the pre-SMA. The NHPs in this study exhibited mild parkinsonian motor signs; additional studies will need to characterize this pathophysiology as the disease evolves to more moderate and severe states while examining the changes that occur across the basal ganglia-thalamocortical network. Our finding of disrupted pre-SMA processing motivates additional investigation into neuromodulation approaches to modify and potentially normalize activity in this region. Previous studies have utilized noninvasive cortical stimulation of the supplementary motor complex in PD patients (typically repetitive transcranial magnetic stimulation, rTMS) with mixed behavioral effects and few specifically targeting pre-SMA (for review see (Rahimpour et al., 2022). Although non-surgical approaches are attractive, it is possible that neuromodulation techniques with finer spatio-temporal precision may be required, for example with chronically implanted electrocorticography (ECoG) electrodes (Morrell and On behalf of the, 2011; Swann et al., 2017), that provide opportunity for more targeted approaches to modify neuronal activity in this area with the goal of improving cognitive-motor processes in PD patients.

## ACKNOWLEDGEMENTS

We would like to thank our colleagues in the Neuromodulation Research Center for helpful comments and critiques related to this study and especially thank our animal core team including Adele DeNicola and Elizabeth McDuell as well as our veterinary and animal care colleagues at the University of Minnesota Research Animal Resources (RAR).

